# Metabolic diversity of microorganisms toward atypical sugar enantiomers

**DOI:** 10.64898/2026.01.17.700050

**Authors:** Hiroaki Nishijima, Kensuke Igarashi, Wataru Kitagawa, Hiro Tabata, Shuji Nakanishi, Souichiro Kato

**Affiliations:** Research Center for Solar Energy Chemistry, Graduate School of Engineering Science, The University of Osaka, Toyonaka, Osaka, Japan; Biomanufacturing Process Research Center, National Institute of Advanced Industrial Science and Technology, Sapporo, Hokkaido, Japan; Presidential Endowed Chair for “Platinum Society”, The University of Tokyo, Bunkyo-ku, Tokyo, Japan; Innovative Catalysis Science Division, Institute for Open and Transdisciplinary Research Initiatives (ICS-OTRI), The University of Osaka, Suita, Osaka, Japan; Department of Biotechnology, Graduate School of Agricultural and Life Sciences, The University of Tokyo, Bunkyo-ku, Tokyo, Japan; Collaborative Research Institute for Innovative Microbiology (CRIIM), The University of Tokyo, Bunkyo-ku, Tokyo, Japan

## Abstract

Life on Earth has long been regarded as homochiral, relying almost exclusively on a single enantiomer of sugars—typically the D-form. However, recent discoveries challenge this paradigm, including the identification of L-glucose-catabolizing bacteria and microbial L-glucoside hydrolases. Despite these findings, the metabolic diversity of organisms toward a broader range of atypical sugar enantiomers and their ecological relevance remains largely unexplored. This study aimed to identify and isolate microorganisms capable of catabolizing atypical enantiomers of diverse sugars. We performed enrichment cultures with either the D- or L-forms of glucose, fructose, xylose, and sorbose, using soil and activated sludge as microbial sources. Microbial growth was observed under all tested conditions, with the dominant taxa varying depending on the sugars supplied. Six phylogenetically distinct bacterial isolates exhibited the ability to catabolize atypical sugar enantiomers, two of which exhibited growth on all tested sugars. These findings uncover a previously unrecognized diversity in microbial sugar metabolisms, providing new insights into the environmental dynamics of atypical sugar enantiomers and offering a novel perspective on the principle of biological homochirality. Furthermore, this work lays a foundation for the development of biomanufacturing processes using racemic sugar mixtures synthesized via abiotic chemical reactions.

## Introduction

In 1890, Emil Fischer achieved the first synthesis of L-glucose (LG) and discovered that it was not fermented by yeast, unlike its mirror-image counterpart D-glucose (DG)^1^. Subsequent studies^2^ demonstrated that LG was not metabolized by rats or *Escherichia coli*, and later, Sun et al.^3^ expanded these findings to include a variety of bacterial and archaeal model species. These foundational investigations established the long-standing consensus that biological systems exhibit homochirality, preferentially utilizing a single enantiomer of sugars, commonly the D-form. Therefore, the origin of homochirality in early life evolution has remained a topic of longstanding debate^4,5^, and the homochirality observed in terrestrial life is widely regarded as a guiding principle in the search for extraterrestrial organisms^6,7^.

Although most sugars utilized by living organisms are in the D-form, several notable exceptions exist. Arabinose, an aldopentose abundant in plant hemicellulose, naturally occurs exclusively in the L-form. Similarly, the deoxy sugars L-fucose (6-deoxy-L-galactose) and L-rhamnose (6-deoxy-L-mannose) are abundantly found as integral constituents of seaweed polysaccharides and other animal-, plant-, and bacteria-derived glycans^8^. Galactose, an aldohexose, is primarily present as the D-enantiomer in nature; however, its L-form is an integral constituent of rhamnogalacturonan-II, a component of plant pectin, making it an exceptional sugar for which both enantiomers abundantly exist in nature^9,10^. For these naturally abundant L-sugars, numerous microorganisms capable of catabolizing them have been identified, and their associated metabolic pathways have been characterized^11,12^.

Furthermore, recent findings have begun to challenge the long-standing paradigm of biological homochirality, with reports of organisms synthesizing or utilizing atypical sugar enantiomers previously considered virtually absent in nature. Shimizu et al.^13^ reported the isolation of multiple soil microorganisms capable of growing on LG. This research group elucidated distinct LG catabolic pathways in two phylogenetically unrelated strains: *Paracoccus laeviglucosivorans* (*α-proteobacteria*)^13^ and *Luteolibacter* sp. (*Verrucomicrobiota*)^14^. Shishiuchi et al.^15^ demonstrated that a metagenome-derived enzyme homologous to L-fucosidase exhibits L-glucosidase activity, with higher substrate specificity toward L-glucosides than toward naturally abundant L-fucosides or L-galactosides. Furthermore, Sasaki et al.^16^ reported selective uptake of LG by human cancer cells, highlighting that its biological relevance extends beyond microbial metabolism. In addition, atypical sugar enantiomers such as LG, L-mannose, L-xylose (LX), and D-fucose have been identified as structural components of bacterial exopolysaccharides and as glycosyl moieties in secondary metabolites produced by plants and microbes^17,18^. Collectively, these findings raise the intriguing possibility that such atypical sugar enantiomers may be more widespread and biologically relevant in nature than previously assumed. Nevertheless, to date, no microorganisms have been reported to catabolize atypical sugar enantiomers other than LG, and the diversity and ecological significance of microbial metabolism of atypical sugar enantiomers remain largely unexplored.

In addition to their ecological and evolutionary significance, microbial metabolisms of atypical sugar enantiomers offer new opportunities for biotechnology, specifically, the effective utilization of racemic sugar mixtures synthesized through chemical processes. In recent years, there has been growing demand for alternative feedstocks for biomanufacturing that do not rely on crop-derived edible biomass^19,20^. One promising approach involves the use of organic compounds produced via abiotic methods, such as inorganic catalysis or electrochemical reactions, which proceed far more rapidly than biological processes^21,22,23^. Among these, sugars synthesized through abiotic processes—particularly via the formose reaction—have attracted increasing attention^24,25^. The formose reaction is a well-known sugar-forming process that involves heating formaldehyde under alkaline conditions in the presence of a base catalyst such as calcium hydroxide. Our group has developed a modified formose reaction that operates under neutral conditions, effectively suppressing undesirable side reactions, and we are currently advancing biomanufacturing technologies that utilize the resulting sugar mixtures^26,27,28^. However, this catalytic reaction yields sugars as racemic mixtures, consisting of equal amounts of D- and L-enantiomers. Therefore, to effectively harness the abiotically synthesized sugars for biomanufacturing, it is essential to employ microorganisms capable of catabolizing atypical sugar enantiomers, and/or to engineer microbial strains with the corresponding metabolic pathways.

In this study, to assess the metabolic diversity of microorganisms toward atypical sugar enantiomers, we conducted enrichment cultures and isolation using either the D- or L-form of glucose, fructose, xylose, or sorbose as the sole carbon and energy source. Analysis of the sugar utilization profiles of the resulting isolates revealed that phylogenetically diverse bacteria are capable of catabolizing atypical sugar enantiomers, with some strains able to catabolize all tested sugars.

## Results and Discussion

### Enrichment cultures of microorganisms on various sugars

To demonstrate the presence of microorganisms capable of catabolizing atypical sugar enantiomers, we conducted enrichment cultures using inorganic media supplemented with either the D- or L-form of glucose (DG and LG), fructose (DF and LF), xylose (DX and LX), or sorbose (DS and LS) as the sole carbon and energy source. Glucose, fructose, and xylose represent naturally abundant aldohexose, ketohexose, and aldopentose, respectively, with their D-forms being the naturally occurring enantiomers. Sorbose, although a minor ketohexose in nature, was included due to its facile formation via the formose reaction^26^. Notably, only LS is found in nature, rendering the D-form an atypical enantiomer. Enrichment cultures were conducted using two environmental samples, namely soil and activated sludge, as microbial sources. Enrichment cultures are named according to the source of their inoculum (‘S’ for soil or ‘A’ for activated sludge) and the type of supplemented sugar; for example, the SLG enrichment refers to the culture derived from soil and supplemented with LG.

Microbial growth was observed under all culture conditions (Supplementary Fig. S1). Although cultures supplemented with atypical sugar enantiomers (especially LG and LF) tended to exhibit slower growth, the final optical density at 600 nm (OD_600_) values were comparable across conditions, suggesting that microbial growth was dependent on the consumption of the supplied sugars. To elucidate differences in microbial species that grew on each sugar, microorganisms were collected from the enrichment cultures after three successive enrichments and subjected to microbial community analysis based on 16S rRNA gene amplicon sequencing. The microbial community compositions obtained from each of the triplicate enrichment cultures and the microbial sources (soil and activated sludge), along with the results of principal component analysis based on the microbial community structures are shown in Supplementary Fig. S2. These analyses clearly demonstrate that the microbial communities after enrichments were markedly different from those of the original inocula, indicating that the enrichment process was effective. Furthermore, the microbial community structures were more strongly influenced by the supplemented sugar species than by differences among the triplicate cultures. Accordingly, subsequent analyses and interpretations were based on the community profiles averaged across the triplicate cultures (Fig. 1).

**Fig. 1.**
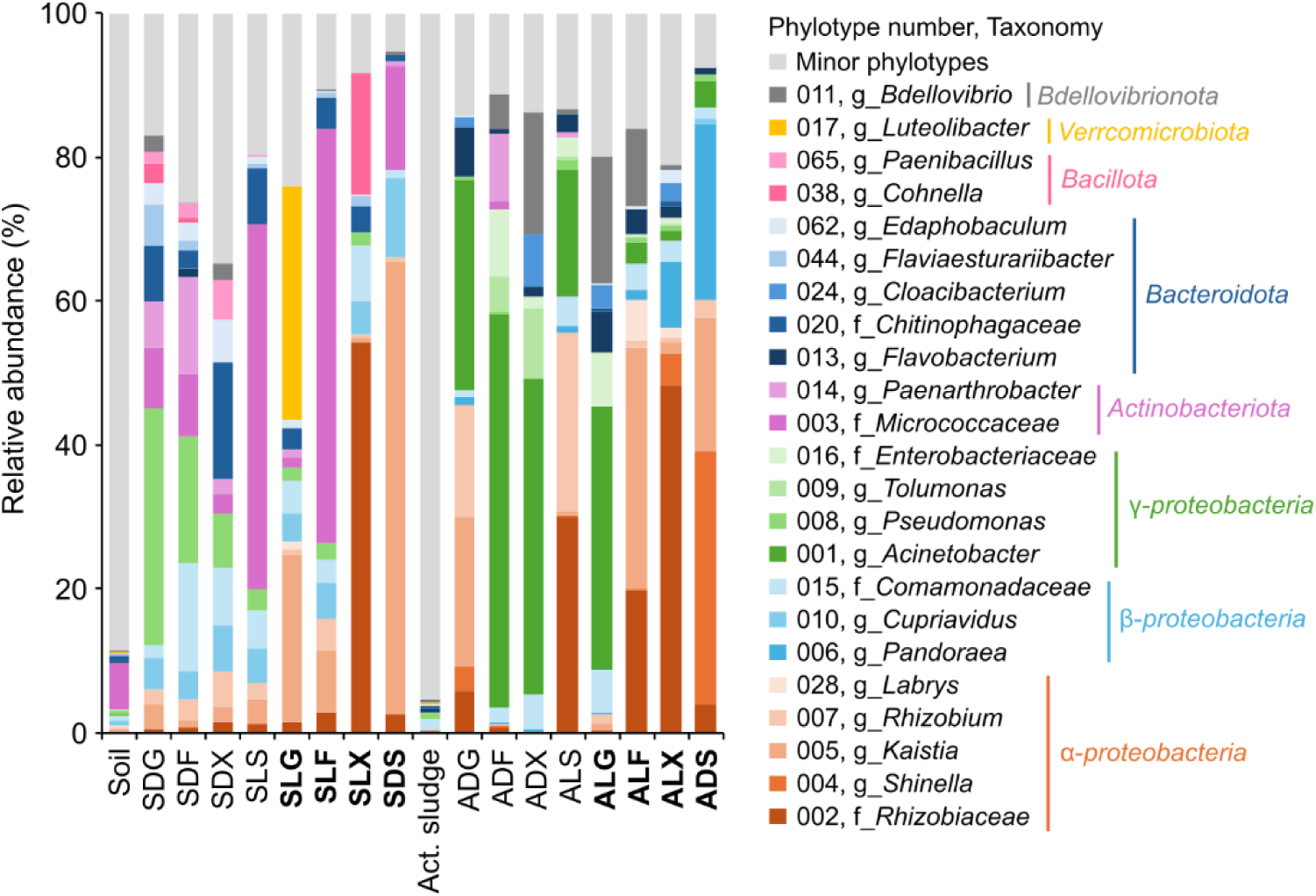
Microbial community profiles of the enrichment cultures with typical or atypical sugar enantiomers. Major phylotypes that accounted for >5% relative abundances in at least one sample were included in the figure, and all others were grouped as minor phylotypes. Enrichment cultures are referred to by a combination of their inoculum source (‘S’ for soil or ‘A’ for activated sludge) and the supplemented sugar. Bold letters indicate enrichment cultures supplemented with atypical sugar enantiomers. Act. sludge; activated sludge.

The enrichment cultures on atypical sugar enantiomers exhibited a tendency toward dominance by *α-proteobacteria* phylotypes. In addition, phylotype 017, which was closely related to *Luteolibacter* (*Verrucomicrobiota*), was dominant only in the SLG enrichment. These results are consistent with previous findings, given that only *P. laeviglucosivorans* (*α-proteobacteria*) and *Luteolibacter* sp. have been identified to date as LG-utilizing bacteria^13,14^.

Furthermore, phylotypes affiliated with *β-* and *γ-proteobacteria*, *Actinomycetota*, *Bacteroidota*, and *Bacillota* were also dominant in the enrichments on atypical sugars, suggesting that the phylogenetic diversity of bacteria capable of utilizing such enantiomers is broader than previously recognized. Several phylotypes were found to be dominant across multiple enrichment cultures supplemented with atypical sugar enantiomers (e.g., phylotypes 002, 003, and 005). For example, phylotype 005 (related to genus *Kaistia*) was dominant across multiple enrichment conditions, namely SLG, SLF, ALF, SDS, and ADS, each differing in both microbial source and supplemented sugar, suggesting that such microorganisms possess the metabolic capacity to utilize a broad range of atypical sugars. It should be noted, however, that not all dominant phylotypes are necessarily sugar-utilizing organisms. For instance, phylotype 011, which was dominant in many enrichment cultures, is closely related to the predatory bacterium *Bdellovibrio* sp., and is likely proliferating by preying on sugar-utilizing microbes rather than catabolizing the supplemented sugars. Therefore, isolation and cultivation-based evaluation are required to demonstrate whether the dominant phylotypes are truly capable of utilizing atypical sugar enantiomers.

### Isolation of microorganisms from the enrichment cultures

Microorganisms were isolated by single-colony isolation using each enrichment as the microbial source and agar-solidified media with the same composition as the corresponding enrichment cultures. From each of the 16 enrichment cultures differing in both microbial source and supplemented sugar, eight colonies were isolated. Isolates sharing >98% sequence identity in partial 16S rRNA gene sequences were considered to represent the same species, and a representative strain from each group was selected for subsequent experiments.

A total of 16 distinct strains were isolated from the enrichment cultures supplemented with atypical sugar enantiomers (Table 1). The majority of these (13 strains) were classified into *α-proteobacteria*, most of which (10 strains) belonged to the order *Hyphomicrobiales*. In addition, two strains were identified as *β-proteobacteria*, and one strain belonged to the phylum *Actinomycetota*. Among these, seven strains closely related to the dominant phylotypes identified in the community analysis of the enrichment cultures were subjected to further culture experiments. The isolates obtained from the enrichments supplemented with typical sugars (Supplementary Table S1) were phylogenetically distinct from those derived from the atypical sugar enrichments. These included two *α-proteobacteria* strains (order *Sphingomonadales*), four *γ-proteobacteria* strains, and two *Actinomycetota* strains. Among them, one strain (strain SDG-1) isolated from the SDG enrichment, and closely related to a dominant phylotype in the same enrichment (phylotype 008), was also subjected to culture experiments.

**Table 1.**
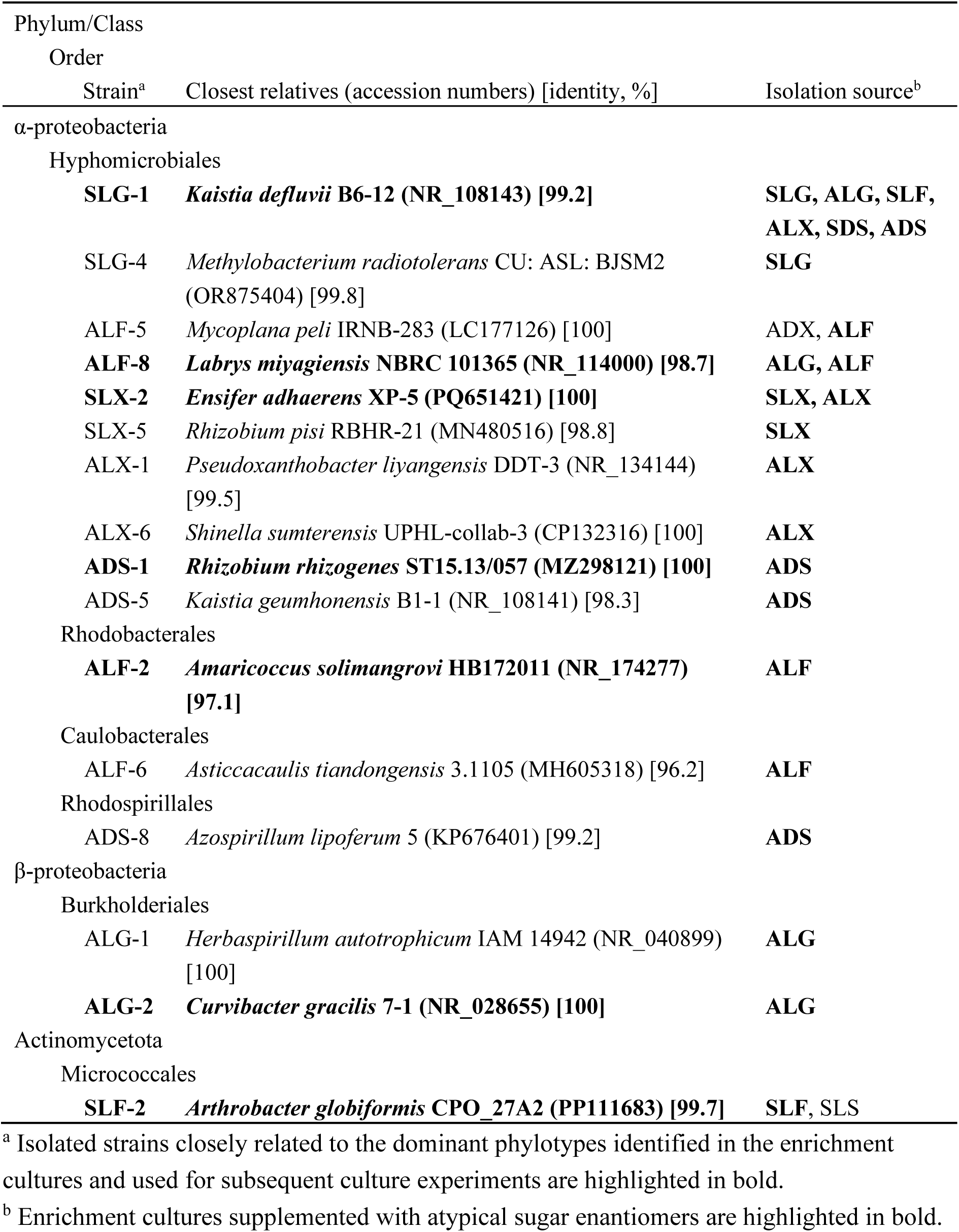
Phylogenetic information of the isolates obtained from enrichment cultures supplemented with atypical sugar enantiomers.

### Sugar utilization properties of isolated strains

To assess the sugar utilization profiles of the isolated strains, growth was evaluated in inorganic liquid media supplemented with individual sugars as the sole carbon and energy source. The model bacterial species, specifically *Escherichia coli* and *Bacillus subtilis,* exhibited growth only on typical sugars (DG, DF, and DX), but not on atypical sugars (Supplementary Figs. S3A, B), consistent with previous findings. It should be noted that slight growth on LF observed in both species was likely due to impurities in the reagent. Among the isolates obtained in this study, strain SDG-1 (99.6% identity to *Pseudomonas putida*), which was isolated from the SDG enrichment and was dominant only in enrichments with typical sugars, grew exclusively on typical sugars (DG and DF) (Supplementary Fig. S3C). Strain ALG-2 (closely related to phylotype 015 and 100% identity to *Curvibacter gracilis*), isolated from the LG enrichment using an LG-supplemented agar medium, showed no growth on any sugar other than DG (Supplementary Fig. S3D). Members of the genus *Curvibacter* are known as oligotrophic bacteria commonly detected in environments such as drinking water^29,30^, suggesting that strain ALG-2 may have relied on organic compounds produced by other microbes in the enrichment cultures or on trace organics present in agar.

In contrast, other six isolated strains exhibited clear growth on at least part of the atypical sugars. One such strain, SLX-2 (closely related to phylotype 012 and 100% identity to *Ensifer adhaerens*), affiliated with the order *Hyphomicrobiales* in the class *α-proteobacteria*, a group frequently observed among the dominant taxa and isolates from atypical sugar enrichments, showed appreciable growth on LX, albeit with a delayed onset compared to growth on typical sugars such as DG, DF, and DX (Fig. 2A). This represents the first experimental evidence of microbial growth on LX. Another *Hyphomicrobiales* strain, ADS-1 (closely related to phylotype 004 and 100% identity to *Rhizobium rhizogenes*), also demonstrated the ability to grow on the atypical sugar DS at a rate comparable to that observed with typical sugars (Fig. 2B). Furthermore, two additional *Hyphomicrobiales* strains, SLG-1 (closely related to phylotype 005 and 99.2% identity to *Kaistia defluvii*) and ALF-8 (closely related to phylotype 028 and 98.7% identity to *Labrys miyagiensis*), exhibited growth on all eight tested sugars, including both typical and atypical enantiomers (Figs. 2C, D). These findings represent the first report of microorganisms capable of catabolizing multiple atypical sugars. In contrast, the model *Hyphomicrobiales* bacterium, *Sinorhizobium meliloti* failed to grow on any of the atypical sugars, while it exhibited appreciable growth on typical sugars (Supplementary Fig. S3E). Collectively, these results demonstrate that the order *Hyphomicrobiales* harbors a diverse array of strains capable of catabolizing atypical sugars, while this metabolic trait is not uniformly distributed across the lineage.

**Fig. 2.**
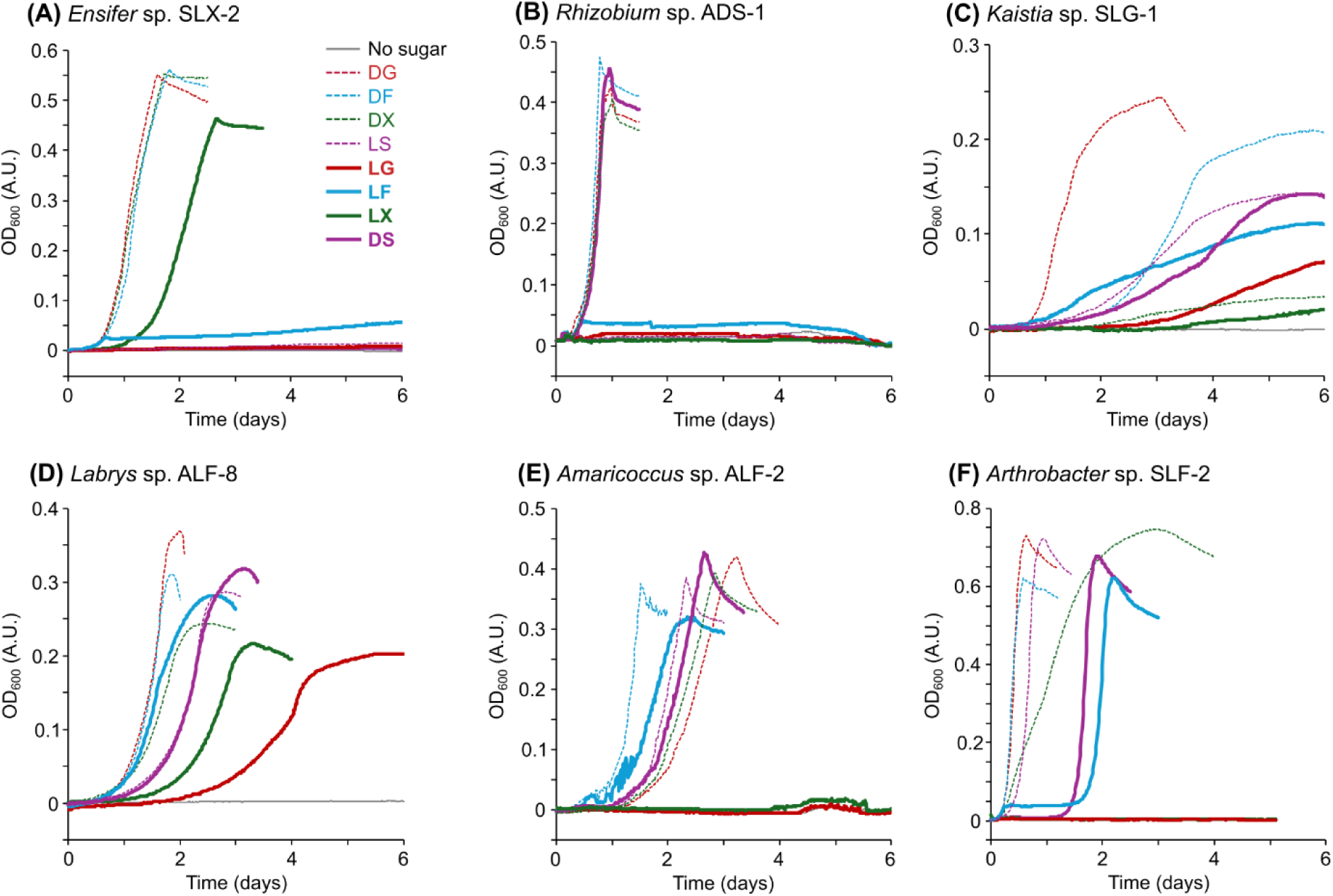
Growth properties of atypical sugar-utilizing isolates. Growth curves on typical sugar enantiomers (D-glucose [DG], D-fructose [DF], D-xylose [DX], and L-sorbose [LS]) are shown as dashed lines, while those on atypical enantiomers (L-glucose [LG], L-fructose [LF], L-xylose [LX], and D-sorbose [DS]) are shown as bold lines. Representative results from five independent cultures are presented. (A) *Ensifer* sp. SLX-2, (B) *Rhizobium* sp. ADS-1, (C) *Kaistia* sp. SLG-1, (D) *Labrys* sp. ALF-8, (E) *Amaricoccus* sp. ALF-2, and (F) *Arthrobacter* sp. SLF-2.

Strain ALF-2 (closely related to phylotype 043 and 97.1% identity to *Amaricoccus solimangrovi*) was isolated from the ALF enrichment culture and classified within the order *Rhodobacterales* of the *α-proteobacteria*, the same order as the known LG-utilizing bacterium *P. laeviglucosivorans*. ALF-2 exhibited growth not only on four typical sugars but also on the atypical sugars LF and DS (Fig. 2E). In contrast, the model *Rhodobacterales* strain, *Paracoccus denitrificans* showed no growth on any of the atypical sugars (Supplementary Fig. S3F). These results suggest that the ability to catabolize atypical sugar enantiomers is not a conserved trait across the order *Rhodobacterales*, as also observed for the order *Hyphomicrobiales*.

Strain SLF-2 (99.7% identity to *Arthrobacter globiformis*), classified within the phylum *Actinomycetota*, was closely related to phylotype 003 that dominated the SLF and SLS enrichment cultures. SLF-2 exhibited the ability to grow on the atypical sugars (LF and DS) as well as four typical sugars (Fig. 2F). These results indicate that, in addition to previously reported atypical sugar-utilizing taxa within the *α-proteobacteria* and *Verrucomicrobiota*, such metabolic potential also exists within the phylum *Actinomycetota*.

### Demonstration of atypical sugar consumption

Although the six atypical sugar-utilizing strains identified in this study tended to grow more slowly on atypical sugar enantiomers compared to their corresponding typical counterparts, the maximum OD_600_ values were generally comparable (Fig. 2). This observation suggests that these bacteria are capable of fully catabolizing atypical sugars and utilizing the resulting energy to support growth. To directly confirm the consumption of atypical sugars, strain ALF-8 was cultured in minimal medium supplemented with either DG, LG, DX, or LX, and sugar concentrations in the medium were monitored throughout the growth period (Fig. 3). For both glucose and xylose, while sugar consumption was slower when the L-enantiomer was provided, LG and LX were ultimately depleted, and the OD_600_ values increased in parallel with sugar consumption. These results clearly demonstrate that atypical sugars are indeed catabolized and utilized as energy and carbon sources.

**Fig. 3.**
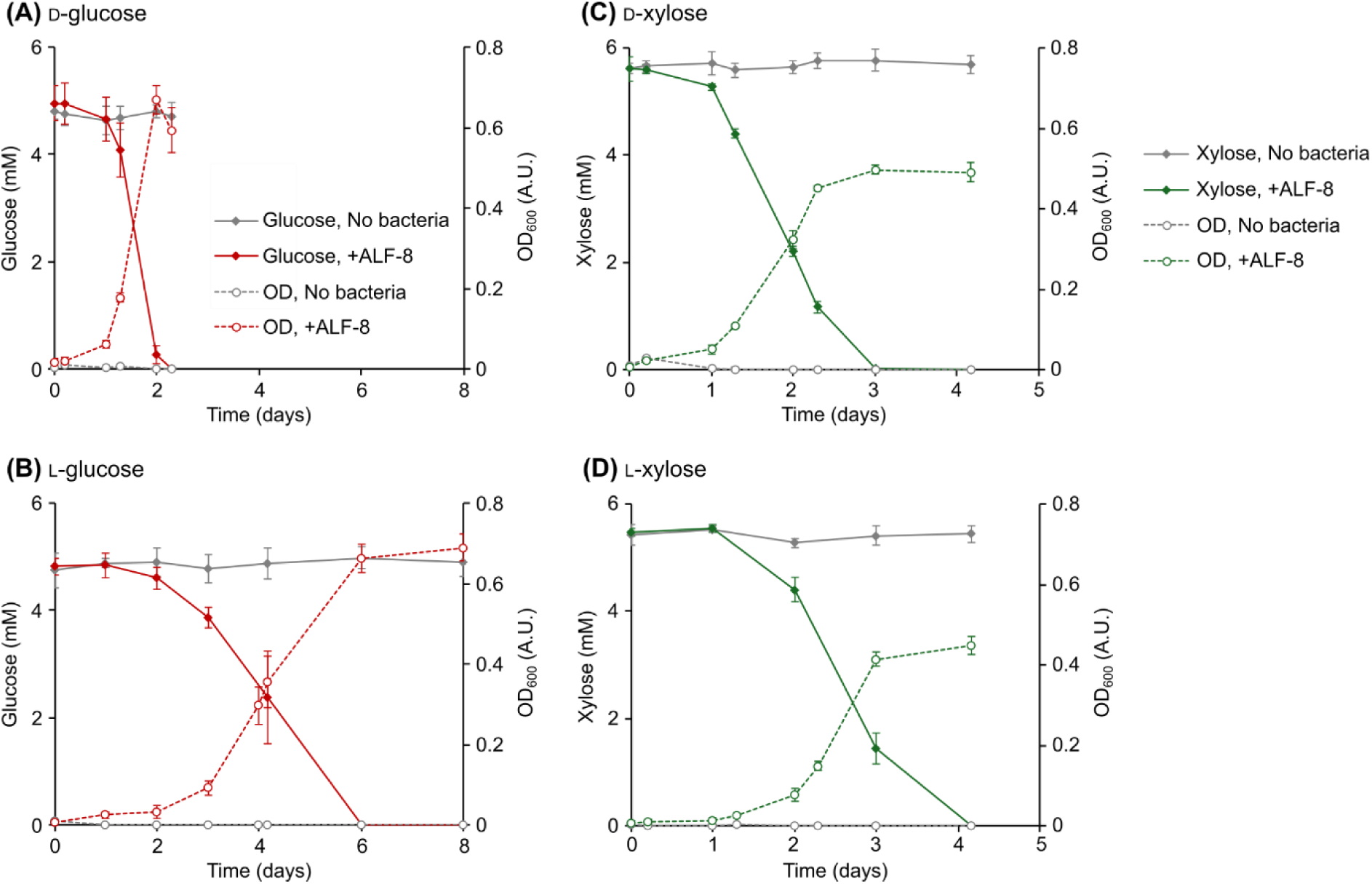
Growth of the atypical sugar-utilizing isolate coupled with sugar consumption. *Labrys* sp. ALF-8 was cultured in inorganic medium supplemented with 1 g/L of D-glucose (A), L-glucose (B), D-xylose (C), or L-xylose (D). Sugar concentrations (solid lines) and bacterial growth (OD_600_, dashed lines) were monitored over time. Medium without bacterial inoculation served as the control. Data represent means of three independent cultures. Error bars indicate standard deviations.

### Implications

In this study, we identified six strains of atypical sugar-utilizing bacteria classified within *α-proteobacteria* and *Actinomycetota*. Together with previously reported LG-utilizing bacteria, our findings indicate that the atypical sugar-utilizing ability is distributed across diverse lineages, including at least the *α-proteobacteria*, *Actinomycetota*, and *Verrucomicrobiota*. Within the *α-proteobacteria*, atypical sugar-utilizing strains were found in two distinct orders, *Hyphomicrobiales* and *Rhodobacterales*, yet model organisms from both orders lacked the ability to utilize atypical sugars. Furthermore, although strains SLX-2 and ADS-1 are phylogenetically very closely related (96.1% 16S rRNA gene identity) within the family *Rhizobiaceae*, they exhibited distinct sugar utilization profiles (Figs. 2A, B). Taken together, these observations suggest that the ability to catabolize atypical sugars is not a broadly conserved trait within specific phylogenetic groups, but rather a feature independently acquired at the family, genus, or species level across multiple lineages.

To date, the only characterized metabolic pathways for atypical sugar enantiomers involve LG metabolisms in *P. laeviglucosivorans*^13^ and *Luteolibacter* sp.^14^. In both cases, LG is metabolized via pathways originally associated with structurally similar typical sugars, scyllo-inositol and L-galactose, respectively, suggesting that LG metabolisms may occur incidentally due to promiscuous enzymatic activity. In contrast, this study identified bacterial strains capable of utilizing multiple atypical sugars (e.g., strains SLG-1 and ALF-8, Figs. 2C, D), as well as strains that exhibited comparable growth rates on atypical and typical sugars despite differences in lag phase duration (e.g., strains ADS-1 and SLF-2, Figs. 2B, F). These characteristics are difficult to explain solely by incidental metabolism through promiscuous enzymes and instead suggest the presence of dedicated enzymes and pathways specialized for the metabolism of each atypical sugar. Further investigation into the metabolic systems of the atypical sugar-utilizing strains identified in this study, including the identification of relevant pathways, evaluation of enzyme specificity, and elucidation of regulatory mechanisms, will be essential to uncover the molecular basis of these unique metabolic capabilities. Furthermore, elucidating the metabolic pathways for diverse atypical sugars could significantly contribute to the development of next-generation biomanufacturing technologies that utilize chemically synthesized racemic sugar mixtures, particularly via the formose reaction^26,27^.

Notably, despite being pre-cultured in DG-supplemented medium prior to the growth assays, strain ALF-2 exhibited the slowest growth on DG, while demonstrating better growth on other sugars, including the atypical sugars LF and DS (Fig. 2E). Most microorganisms preferentially utilize carbon sources that are more readily available and support faster growth, generally DG, through a mechanism known as carbon catabolite repression (CCR)^31^. Indeed, all other isolates and model strains examined in this study showed their fastest growth on DG (Fig. 2 and Supplementary Fig. S3). To date, several microorganisms have been reported to exhibit a phenomenon known as reverse CCR, in which carbon substrates other than DG are preferentially utilized. For example, members of the genera *Streptomyces* and *Bifidobacterium* preferentially catabolize certain oligosaccharides over DG^32,33^, while *Pseudomonas* species have been shown to favor organic acids over sugars^34^. In addition, recent studies have demonstrated that the marine bacterium *Phaeobacter inhibens* preferentially utilizes mannitol and N-acetylglucosamine over glucose^35^. However, the preferential utilization of sugars that are scarcely found in nature—let alone atypical sugar enantiomers—has not been previously documented. Although further investigation is required to substantiate this observation, our findings raise the possibility that certain atypical sugar-utilizing bacteria, such as strain ALF-2, may possess previously unrecognized carbon substrate preferences.

In addition to sugars, amino acids represent another class of biomolecules that exhibit homochirality in extant life, with L-enantiomers serving as the typical form. The prevalence of L-amino acids is thought to be linked to the stereospecific affinity of RNA—composed of D-ribose—for L-amino acids^36,37^. While it remains unclear whether the homochirality of RNA led to the selection of L-amino acids, or vice versa^38^, it is evident that D-sugars and L-amino acids constitute the dominant enantiomeric forms utilized by life on Earth. Nevertheless, numerous examples of D-amino acid utilization in biological systems have been documented^39^. A prominent example is the incorporation of D-amino acids into peptidoglycan of bacterial cell walls. This usage is thought to have evolved as a strategy to evade degradation by conventional proteases^40^. A similar rationale may apply to sugars. L-galactose, an abundant L-sugar found in plant and algal polysaccharides^9,41^, as well as atypical sugar enantiomers such as LG, LX, L-mannose, and D-fucose, identified as structural components of bacterial exopolysaccharides and as glycosyl moieties in various secondary metabolites^17,18^, may have been adopted to resist enzymatic degradation. It is conceivable that numerous atypical sugars, though not yet identified, may be utilized by a broader range of organisms. As demonstrated in this study, the presence of atypical sugar-catabolizing capabilities across diverse microbial lineages supports this hypothesis. Continued exploration of naturally occurring atypical sugars and the microorganisms that metabolize them may reveal previously unrecognized ecological significance of these structurally uncommon carbohydrates. Furthermore, insights into enzymes acting on atypical sugar enantiomers may provide valuable knowledge and resources for the emerging field of synthetic biology for ‘mirror life’, organisms composed of L-sugars and D-amino acids^42,43,44^.

## Conclusion

In this study, we identified six phylogenetically distinct bacterial strains capable of utilizing atypical sugar enantiomers through enrichment and isolation using both D- and L-forms of four different sugars. Notably, these included previously unreported LF-, LX-, and DS-assimilating strains, as well as microbes capable of metabolizing multiple atypical sugar enantiomers. Our findings provide the first evidence that microorganisms possess diverse capabilities for utilizing atypical sugar enantiomers, suggesting that such sugars may be more universally accessible to biological systems than previously assumed.

## Materials and methods

### Bacterial strains and culture conditions

*E. coli* strain K12 (ATCC 12435) was routinely cultured in the M9 minimal medium^45^ supplemented with 1 g/L DG. *B. subtilis* strain 168 (JCM 10629), *S. meliloti* JCM 20682^T^, and *P. denitrificans* JCM 21484^T^ were regularly cultured in the IET minimal medium^28^ supplemented with 1 g/L of DG. Stock solutions of each sugar were prepared by filter sterilization using 0.22-μm pore filters (MilliporeSigma, Burlington, MA, US) and subsequently added to the autoclaved media. Growth of bacteria was measured by monitoring the OD_600_ of the culture solution with a 96-well plate reader (BioTek LogPhase 600, Agilent Technologies, Santa Clara, USA) with the uninoculated medium as the control. Sugars in the supernatant of culture solution were quantified by using a high-performance liquid chromatography (HPLC) system (D-2000 LaChrom Elite HPLC system; Hitachi, Tokyo, Japan) equipped with Aminex HPX-87H column (300 mm, 7.8 mm I.D.; Bio-Rad Laboratories, Hercules, CA, United States), and a refractive index detector (L-2490, Hitachi, Tokyo, Japan), as described previously^46^.

### Enrichment cultures and isolation of sugar-catabolizing microorganisms

Enrichments of sugar-catabolizing microorganisms were performed using the IET minimal medium supplemented with 1 g/L of DG, LG, DF, LF, DX, LX, DS or LS. Approximately 50 μL of suspension of woodland soil in The University of Osaka (Osaka, Japan) or activated sludge for treating municipal wastewater (Sapporo, Japan) was inoculated as a microbial source.

Enrichment cultures were incubated at 30°C with agitation (180 rpm) until visible growth was confirmed. After three successive enrichment cultures, sugar-catabolizing microorganisms were isolated by single colony isolation using the agar-solidified IET minimal medium supplemented with 1 g/L of each sugar used for the enrichment cultures. The isolates were further purified by single colony isolation on the same medium at least twice. All isolates were routinely cultured in the IET minimal medium supplemented with 1 g/L DG at 30°C with agitation (180 rpm).

Partial 16S rRNA gene sequences of the isolates were determined by the direct sequencing of PCR products with the PCR primer pair 27F/1492R and sequencing primer 533R as described previously^47^. Isolates with >98% 16S rRNA gene sequence identities were designated as the same phylotypes. The closest relatives of the isolates were inferred using the BLAST program^48^. For the eight isolated strains that were dominant in the enrichment cultures, almost full-length 16S rRNA genes were determined by the direct sequencing of PCR products with the primer pair 27F/1492R as described previously^47^ and subjected to further experiments. These eight strains have been deposited in the Japan Collection of Microorganisms (JCM) as follows: *Arthrobacter* sp. SLF-2 (JCM 38031), *Rhizobium* sp. ADS-1 (JCM 38037), *Amaricoccus* sp. ALF-2 (JCM 38038), *Labrys* sp. ALF-8 (JCM 38039), *Curvibacter* sp. ALG-2 (JCM 38040), *Pseudomonas* sp. SDG-1 (JCM 38044), *Kaistia* sp. SLG-1 (JCM 38045), and *Ensifer* sp. SLX-2 (JCM 38046).

### Microbial community analysis

DNA was extracted using FAST DNA Spin Kit for Soil (MP Biomedicals, Irvine, US) and then purified using QIAquick PCR Purification Kit (Qiagen, Hilden, Germany) following the manufacturers’ instructions. PCR amplification of bacterial and archaeal 16S rRNA gene V4 region was performed using the purified DNA as a template, KAPA HiFi HotStart ReadyMix (Roche Sequencing Solutions, Pleasanton, CA, US), and primers 515’F/806R^49^. The thermal conditions were as follows: initial thermal denaturation at 95°C for 3 min, followed by 25 cycles of heat denaturation at 95°C for 30 sec, annealing at 55°C for 30 sec and extension at 72°C for 30 sec; and a final cycle at 72°C for 5 min. The PCR product was separated on agarose gel electrophoresis and then extracted by gel extraction using QIAquick PCR purification kit (Qiagen). The cleaned PCR product was used as the template for the subsequent index PCR reaction to incorporate a dual-index Nextera barcode and the remaining Illumina adapter sequence (Illumina, San Diego, CA, USA). The PCR condition was the same as above, but the amplification cycle was limited to 8 cycles. The resulting index PCR product was subjected to agarose gel electrophoresis to confirm length of amplified fragments and purified as above. Each purified PCR product was quantified using the Qubit 4 and its dsDNA BR Assay Kit (Thermo Fisher Scientific, Waltham, MA, USA) and then mixed to prepare amplicon DNA library. The library was subjected to sequencing analysis with an iSeq 100 sequencing system according to the manufacturer’s instructions, utilizing an Illumina iSeq™ 100 i1 Reagent v2 kit (Illumina). The bioinformatics analysis was performed using the QIIME 2 pipeline v.2024.10 ^50^. The raw sequence data (FASTQ) were imported into QIIME 2. Then, data were demultiplexed, quality filtering was performed using the q2-demux plugin, and sequence denoising was carried out using DADA2. The taxonomic classification of the amplicon sequence variants (ASVs) was performed using the classify-consensus-blast classifier of the q2-feature-classifier plugin against the SILVA v.138.99 database for 16S ASVs (https://docs.qiime2.org/2024.10/data-resources/). Principal component analysis was performed using Principal Component Analysis Calculator available at Statistics Kingdom (https://www.statskingdom.com/pca-calculator.html).

## Acknowledgements

The authors would like to express appreciation to Ms. Ai Miura, Ms. Mika Yamamoto, Ms. Keiko Fujikawa and Ms. Sharbanee Mitra for technical assistance. The work was partly supported by the JST-Mirai Program Grant No. JPMJMI22E5, Japan.

## Author contributions

Conceptualization, funding acquisition, and data curation, S.N. and S.K.; methodology, resources, visualization, and writing–original draft preparation, H.N. and S.K.; investigation, formal analysis, and validation; H.N., K.I., W.K., and H.T.; and writing–review and editing, H.N., K.I., W.K., H.T., S.N., and S.K.

## Data availability statement

The nucleotide sequence data obtained from the isolates in the present study are available in the DDBJ/EBI/NCBI databases under accession numbers LC903028 to LC903035 and LC903146 to LC903161. Any additional information required to reanalyze the data reported in this paper is available from the lead contact (Souichiro Kato) upon request.

## Additional Information

The authors declare no competing interests.

