## Supplementary Fig S1, S2, S3 and Table S1 for "Metabolic diversity of microorganisms toward atypical sugar enantiomers"

**Supplementary information**

Supplementary Figs. S1, S2, S3

Supplementary Table S1

**Supplementary Fig. S1.** Growth curves of the enrichment cultures with typical or atypical sugar enantiomers using soil (A-H) or activated sludge (I-P) as the microbial inocula, respectively. Enrichment cultures are named according to the source of their inoculum ('S' for soil or 'A' for activated sludge) and the type of supplemented sugar, i.e., typical sugar enantiomers (D-glucose [DG], D-fructose [DF], D-xylose [DX], or L-sorbose [LS]) or atypical enantiomers (L-glucose [LG], L-fructose [LF], L-xylose [LX], or D-sorbose [DS]). All data from three independent enrichment cultures are shown.

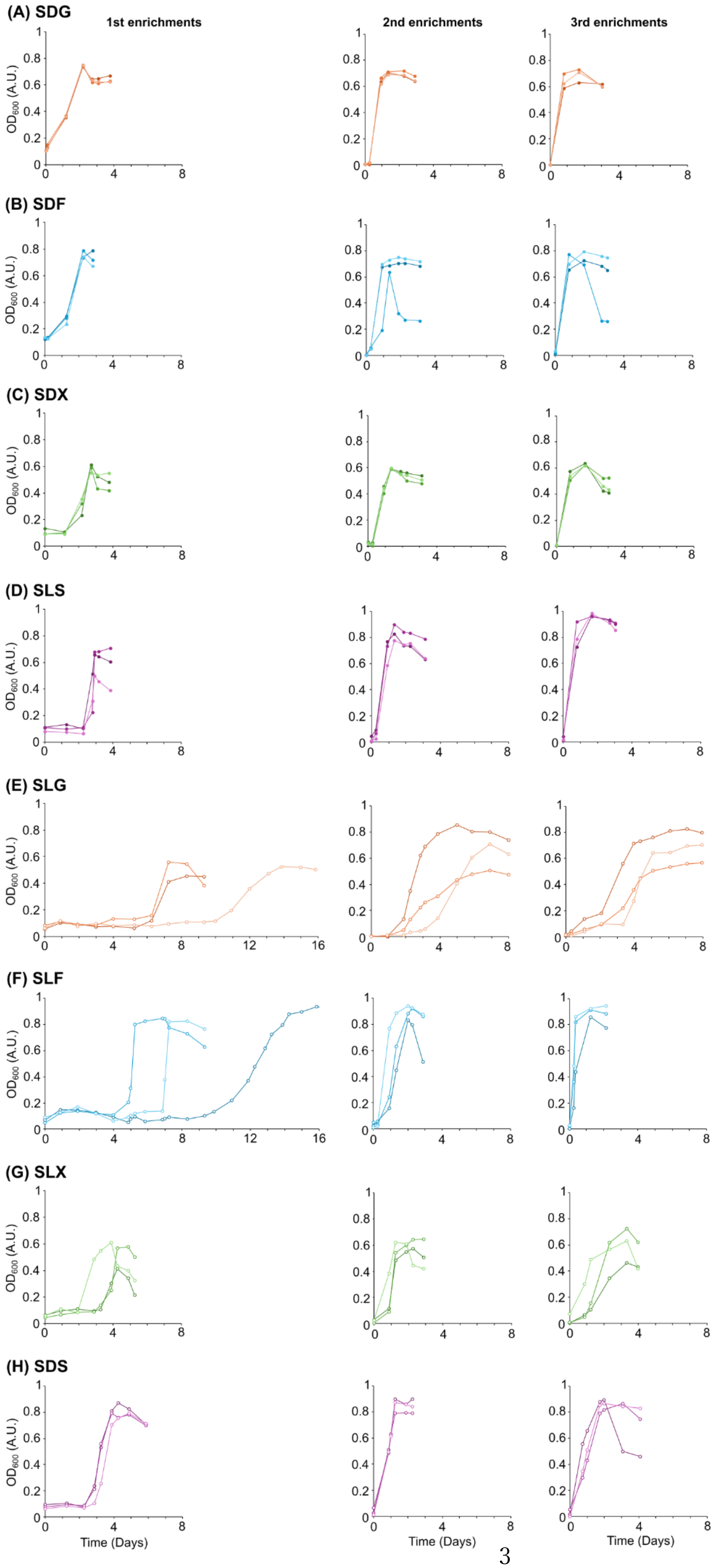
(A) SDG
(B) SDF
(C) SDX
(D) SLS
(E) SLG
(F) SLF
(G) SLX
(H) SDS
1st enrichments
2nd enrichments
3rd enrichments
OD600 (A.U.)
Time (Days)

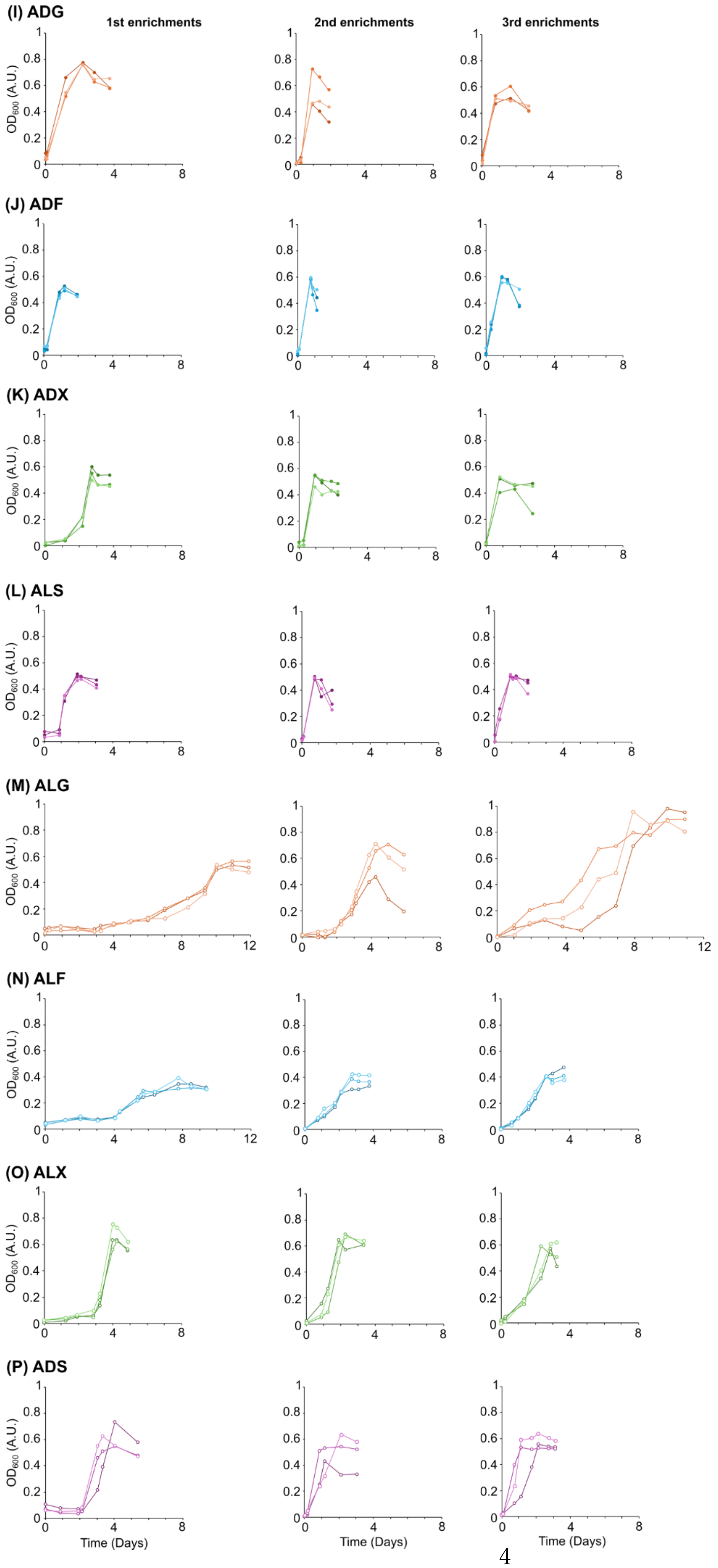
(I) ADG
1st enrichments
2nd enrichments
3rd enrichments
(J) ADF
(K) ADX
(L) ALS
(M) ALG
(N) ALF
(O) ALX
(P) ADS
$OD_{600}$ (A.U.)
Time (Days)

**Supplementary Fig. S2.** Microbial community profiles (A, B) and their principal component analysis (C, D) of the enrichment cultures with typical or atypical sugar enantiomers using soil (A, C) or activated sludge (Act. sludge, B, D) as the microbial inocula, respectively. Major phylotypes that accounted for >5% relative abundances in at least one sample were included in the figure, and all others were grouped as minor phylotypes. Enrichment cultures are named according to the source of their inoculum (‘S’ for soil or ‘A’ for activated sludge) and the type of supplemented sugar. Bold letters indicate enrichment cultures supplemented with atypical sugar enantiomers. -a, -b, and -c indicate triplicate cultures. Note that the ALF-b and -c enrichment cultures were excluded from the analysis due to insufficient PCR product obtained from two of the three replicates.

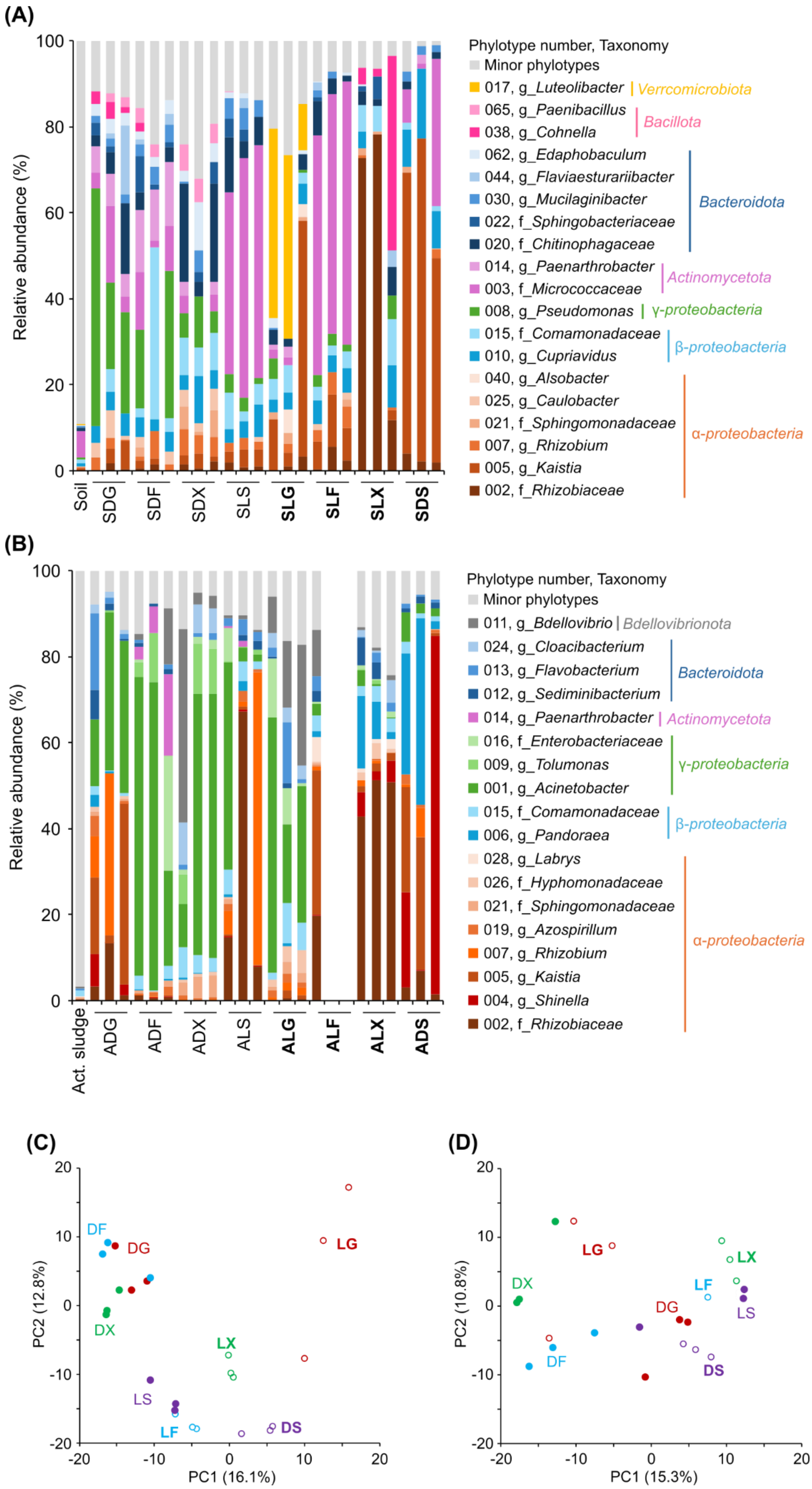

(A)
Relative abundance (%)
Soil
SDG
SDF
SDX
SLS
SLG
SLF
SLX
SDS
Phylotype number, Taxonomy
Minor phylotypes
017, g_Luteolibacter
Verrcomicrobiota
065, g_Paenibacillus
038, g_Cohnella
Bacillota
062, g_Edaphobaculum
044, g_Flaviaesturariibacter
030, g_Mucilaginibacter
022, f_Sphingobacteriaceae
020, f_Chitinophagaceae
Bacteroidota
014, g_Paenarthrobacter
003, f_Micrococcaceae
Actinomycetota
008, g_Pseudomonas
γ-proteobacteria
015, f_Comamonadaceae
010, g_Cupriavidus
β-proteobacteria
040, g_Alsobacter
025, g_Caulobacter
021, f_Sphingomonadaceae
007, g_Rhizobium
005, g_Kaistia
002, f_Rhizobiaceae
α-proteobacteria
(B)
Relative abundance (%)
Act. sludge
ADG
ADF
ADX
ALS
ALG
ALF
ALX
ADS
Phylotype number, Taxonomy
Minor phylotypes
011, g_Bdellovibrio
Bdellovibrionota
024, g_Cloacibacterium
013, g_Flavobacterium
012, g_Sediminibacterium
Bacteroidota
014, g_Paenarthrobacter
Actinomycetota
016, f_Enterobacteriaceae
009, g_Tolumonas
001, g_Acinetobacter
γ-proteobacteria
015, f_Comamonadaceae
006, g_Pandoraea
β-proteobacteria
028, g_Labrys
026, f_Hyphomonadaceae
021, f_Sphingomonadaceae
019, g_Azospirillum
007, g_Rhizobium
005, g_Kaistia
004, g_Shinella
002, f_Rhizobiaceae
α-proteobacteria
(C)
PC2 (12.8%)
PC1 (16.1%)
DF
DG
DX
LG
LX
LS
LF
DS
(D)
PC2 (10.8%)
PC1 (15.3%)
LG
DX
LX
LF
DG
LS
DF
DS

**Supplementary Fig. S3.** Growth properties of model microorganisms and isolates without atypical sugar-utilizing abilities. Growth curves on typical sugar enantiomers (D-glucose [DG], D-fructose [DF], D-xylose [DX], and L-sorbose [LS]) are shown as dashed lines, while those on atypical enantiomers (L-glucose [LG], L-fructose [LF], L-xylose [LX], and D-sorbose [DS]) are shown as bold lines. Representative results from five independent cultures are presented. (A) *Escherichia coli* strain K12, (B) *Bacillus subtilis* strain 168, (C) *Pseudomonas* sp. SDG-1, (D) *Curvibacter* sp. ALG-2, (E) *Sinorhizobium meliloti* JCM 20682$^T$, and (F) *Paracoccus denitrificans* JCM 21484$^T$.

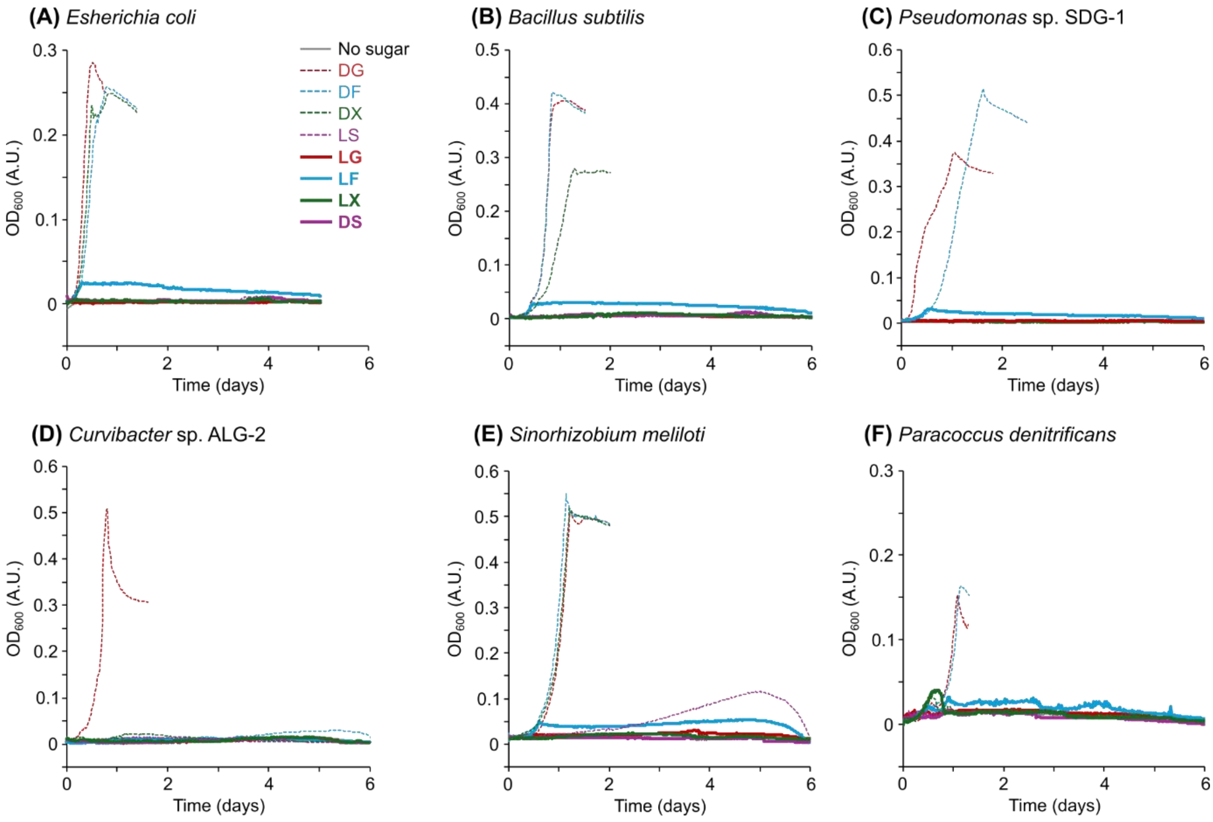

**Supplementary Table S1.** Phylogenetic information of the isolates obtained from enrichment cultures supplemented with typical sugar enantiomers.

| Phylum/Class<br>Order<br>Strain[a] | Closest relatives (accession numbers) [identity, %] | Isolation source[b] |
|---|---|---|
| α-proteobacteria | | |
| Sphingomonadales | | |
| SDX-1 | *Novosphingobium silvae* CRRU10 (OQ195983) [99.0] | ADG, SDX, ADX |
| SDX-8 | *Sphingobium yanoikuyae* KUDC1818 (KC355325) [99.8] | SDX |
| γ-proteobacteria | | |
| Pseudomonadales | | |
| **SDG-1** | ***Pseudomonas putida* IHB B 1071 (KF475834) [99.6]** | SDG |
| Aeromonadales | | |
| ADF-1 | *Tolumonas auensis* DSM 9187 (NR_074805) [98.9] | ADG, ADF |
| Enterobacterales | | |
| ALS-5 | *Klebsiella variicola* FDAARGOS_628 (CP050958) [100] | ADG, ADF, ALS |
| Moraxellales | | |
| ALS-6 | *Acinetobacter brisouii* ABIP 1234 (OQ061524) [99.6] | ADX, ALS |
| Actinomycetota | | |
| Micrococcales | | |
| SDG-4 | *Pseudarthrobacter phenanthrenivorans* H31 (KC934818) [99.1] | SDG, SDF |
| SLS-1 | *Pseudarthrobacter defluvii* CNMUM20.6 (MN004832) [100] | SLS |

[a] Isolated strains used for subsequent culture experiments are highlighted in bold.

[b] Enrichment cultures supplemented with atypical sugar enantiomers are highlighted in bold.
